# Cooperative regulation of C1-domain membrane recruitment polarizes atypical Protein Kinase C

**DOI:** 10.1101/2022.01.01.474707

**Authors:** Kimberly A. Jones, Michael L. Drummond, Rhiannon R. Penkert, Kenneth E. Prehoda

## Abstract

Recruitment of the Par complex protein atypical Protein Kinase C (aPKC) to a specific membrane domain is a key step in the polarization of animal cells. While numerous proteins and phospholipids interact with aPKC, how these interactions cooperate to control its membrane recruitment has been unknown. Here we identify aPKC’s C1 domain as a phospholipid interaction module that targets aPKC to the membrane of *Drosophila* neural stem cells (NSCs). The isolated C1 binds the NSC membrane in an unpolarized manner during interphase and mitosis and is uniquely sufficient among aPKC domains for targeting. Other domains, including the catalytic module and those that bind the upstream regulators Par-6 and Bazooka, restrict C1’s membrane targeting activity – spatially and temporally – to the apical NSC membrane during mitosis. Our results suggest that aPKC polarity results from cooperative activation of autoinhibited C1-mediated membrane binding activity.

## Introduction

The Par complex polarizes animal cells by excluding specific cortical factors from the Par cortical domain (Lang and Munro, 2017; Venkei and Yamashita, 2018). During polarization the proteins Par-6 and atypical Protein Kinase C (aPKC), which make up the complex, are recruited to a specific, continuous region of the cell membrane, such as the apical surface of epithelia (Tepass, 2012), the anterior hemisphere of the *C. elegans* zygote (Nance and Zallen, 2011), or the apical hemisphere of *Drosophila* neural stem cells (NSCs) (Prehoda, 2009). Factors that are directly polarized by the Par complex, such as Miranda and Numb, are aPKC substrates (Atwood and Prehoda, 2009; Smith et al., 2007). Phosphorylation is coupled to removal from the Par domain, causing these substrates to localize to a complementary membrane domain (Bailey and Prehoda, 2015). Thus, the pattern of Par-polarized factors is ultimately determined by the mechanisms that specify aPKC’s membrane recruitment and activation.

Many aPKC interactions with proteins and phospholipids have been identified (Figure 1A), although how they collaborate to polarize aPKC remains poorly understood. Bazooka (Baz aka Par-3) and Par-6 form direct physical contacts with aPKC, and each protein has possible pathways for membrane recruitment: Baz through direct interactions with the membrane and Par-6 through interactions with prenylated Cdc42 (Joberty et al., 2000; Lin et al., 2000; Krahn et al., 2010). Several direct interactions with phospholipids have also been reported, including with ceramide (Wang et al., 2005), sphingosine-1-phosphate (Kajimoto et al., 2019), and phosphoinositides (Dong et al., 2020; Standaert et al., 1997). The aPKC catalytic domain may also play a role in membrane recruitment as perturbations in this domain can cause aPKC to become depolarized (Rodriguez et al., 2017; Hannaford et al., 2019).

**Figure 1.**
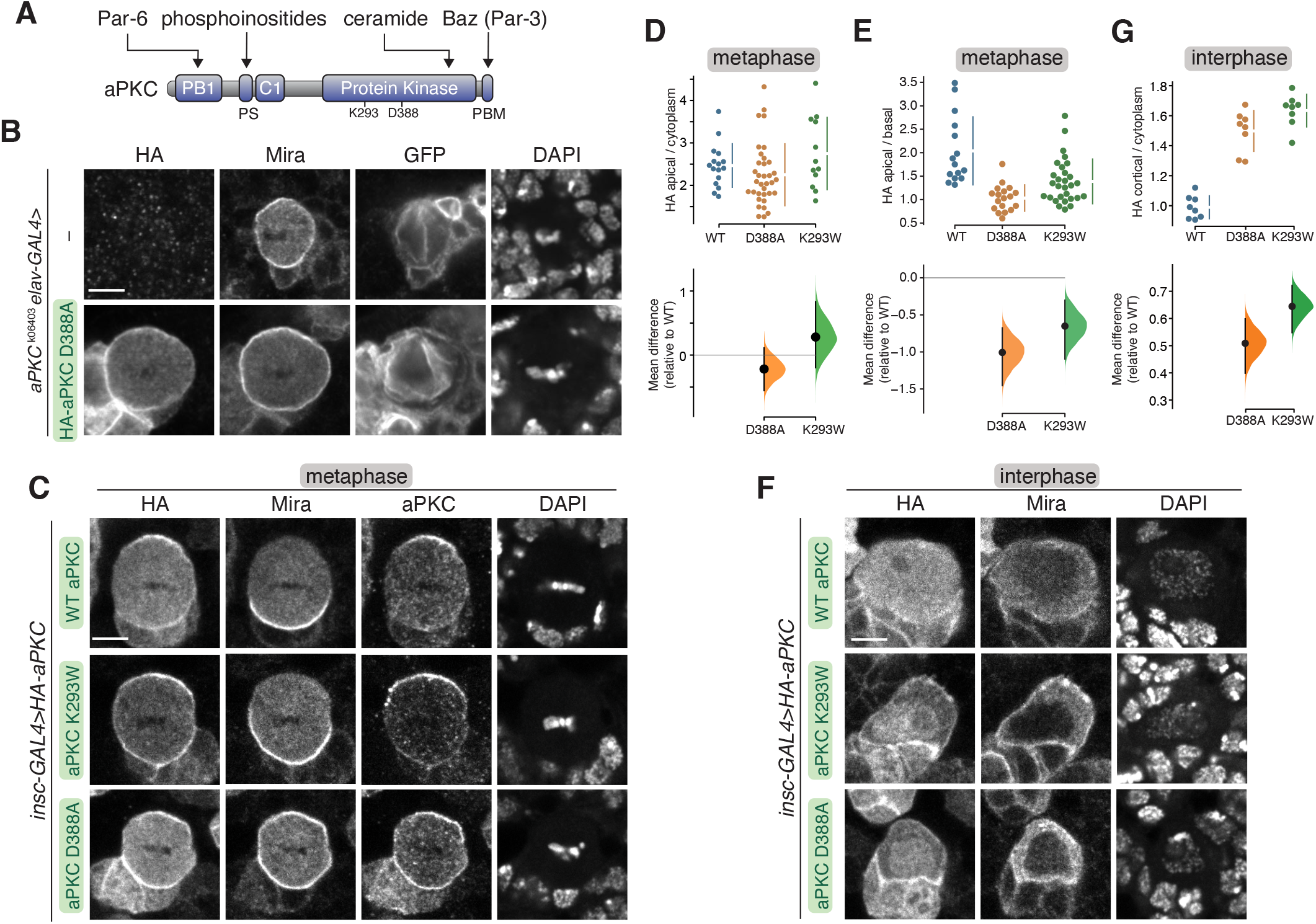
Localization of aPKC with kinase inactivating mutations in larval brain NSCs. **(A)** Domain structure of aPKC showing the location of PB1, PS (pseudosubstrate), C1, kinase domain, PBM (PDZ binding motif), along with location of K293 and D388 residues. **(B)** Localization of HA tagged aPKC harboring the D388A kinase inactivating mutation in a metaphase, positively marked (mCD8-GFP) *aPKC*^k06403^ mutant larval brain NSC with an *aPKC*^K06403^ mutant larval brain NSC shown for comparison. Nucleic acids are shown with DAPI. Scale bar is 5 μm in all panels. **(C)** Localization of HA tagged aPKC harboring either the D388A or K293W kinase inactivation mutations in metaphase larval brain NSCs with endogenous aPKC. The basal cortical marker Miranda, total aPKC (“aPKC”, endogenous and exogenously expressed), and nucleic acid (DAPI) are also shown. **(D,E)** Gardner-Altman estimation plots of the effect of the D388A and K293W mutations on metaphase aPKC membrane recruitment. Apical cortical to cytoplasmic (D) and apical/basal (ED) signal intensities of anti-HA signals are shown for individual metaphase NSCs expressing either HA-WT or HA-D388A or HA-K293W aPKC. Error bar in upper graph represents one standard deviation (gap is mean); error bar in lower graph represents bootstrap 95% confidence interval. **(F)** Localization of HA tagged aPKC harboring either the D388A or K293W kinase inactivation mutations in interphase larval brain NSCs with endogenous aPKC. The basal cortical marker Miranda and nucleic acid (DAPI) are also shown. **(G)** Gardner-Altman estimation plot of the effect of the D388A and K293W mutations on interphase aPKC membrane recruitment. Cortical to cytoplasmic cortical signal intensities of anti-HA signals are shown for individual metaphase NSCs expressing either HA-WT or HA-D388A or HA-K293W aPKC. Error bar in upper graph represents one standard deviation (gap is mean); error bar in lower graph represents bootstrap 95% confidence interval.

While interactions that could potentially recruit aPKC to the membrane have been identified, what is missing is an understanding of how the interactions function together to polarize aPKC at the proper time. One possibility is an avidity model where each interaction is weak, unable to recruit aPKC to the membrane on its own, but the energy provided by multiple interactions allows for recruitment. Alternately, one or more interactions could be sufficient for recruitment but somehow regulated to ensure that targeting only occurs when the appropriate cues are present. These models could be distinguished by determining if any interactions are sufficient for targeting.

We used *Drosophila* NSCs to investigate how aPKC is recruited to the membrane during polarization (Homem and Knoblich, 2012). During interphase, aPKC is cytoplasmic in NSCs but becomes targeted to the apical hemisphere early in mitosis, ultimately concentrating near the apical pole before depolarizing and returning to the cytoplasm as division completes (Oon and Prehoda, 2019). The highly dynamic nature of the NSC polarity cycle makes it possible to assess both spatial (polarized, depolarized, or cytoplasmic) and temporal (interphase or mitotic) aspects of aPKC membrane recruitment regulation.

## Results

### Mutations that inactivate catalytic activity depolarize aPKC

We began our examination of aPKC membrane targeting mechanisms in NSCs by evaluating the role of the catalytic domain. At metaphase, aPKC is highly enriched at the apical membrane of larval brain NSCs (Rolls et al., 2003). We examined the effect of mutations that inactivate aPKC’s catalytic activity on its localization in cells that lacked endogenous aPKC (*apkc*^K06403^ in positively marked clones; Lee and Luo, 1999). Besides aPKC’s localization, we also examined its activity in these metaphase NSCs by determining the localization of Miranda (Mira), an aPKC substrate that is normally restricted to the basal cortex by apical aPKC activity (Figure 1B) (Atwood and Prehoda, 2009; Ikeshima-Kataoka et al., 1997). Consistent with previous observations, we found that in metaphase *apkc*^K06403^ NSCs, Mira was allowed to enter the apical cortex becoming uniformly cortical (Figure 1B). Expression of wild-type aPKC restores the apical aPKC and basal Mira localization found in normally functioning NSCs to positively marked *aPKC^K06403^* metaphase null clones (Holly et al., 2020).

To examine the effect of perturbing catalytic activity, we expressed aPKC harboring a mutation (D388A) that does not have detectable activity in an *in vitro* protein kinase assay (Holly et al., 2019). This mutation is in a residue that coordinates the γ-phosphate of ATP and is thought to allow ATP to bind but prevent phosphotransfer (Cameron et al., 2009). Unlike wild-type aPKC, which is restricted to the apical domain at metaphase, we found that aPKC D388A was largely depolarized, localizing along the entire cortex of positively marked *aPKC*^K06403^ metaphase null clones (Figure 1B). Mira also localized uniformly to the cortex in these cells, confirming that aPKC D388A is inactive both *in vitro* (Holly et al., 2019) and *in vivo* (Figure 1B). We conclude that inactivation of the aPKC catalytic domain depolarizes aPKC in metaphase NSCs.

### Kinase inactive aPKC is not polarized by endogenous aPKC

Our results indicate that mutations that perturb aPKC’s catalytic activity also influence its localization. Previous examinations of chemically inhibited aPKC also found that perturbing catalytic activity caused aPKC to localize to the membrane but in a depolarized manner (Rodriguez et al., 2017; Hannaford et al., 2019). The depolarization caused by perturbations to the kinase domain could be explained if aPKC’s catalytic activity directly participated in its own localization (e.g. by a feedback mechanism). Alternately, perturbations in the kinase domain could alter other aPKC functions. Besides catalyzing phosphotransfer, the aPKC kinase domain also binds a pseudosubstrate in its NH_2_-terminal region causing autoinhibition (Graybill et al., 2012). Mutations or small molecules that influence the active site could perturb the intramolecular interaction in addition to inhibiting catalytic activity. To determine whether the loss of aPKC’s catalytic activity is responsible for the localization defects of aPKC D388A, we examined whether the presence of wild-type, endogenous aPKC with its normal level of catalytic activity could restore aPKC D388A polarity. We also tested the localization of a well-characterized kinase inactive mutation K293W, which blocks ATP binding (Graybill et al, 2012). In NSCs containing endogenous aPKC, both aPKC D388A and K293W remained enriched at the membrane (Figure 1C,D) but depolarized (Figure 1C,E), indicating that aPKC catalytic activity is not sufficient to restore polarity to these proteins. Interestingly, in cells expressing aPKC K293W Mira was basally polarized but in cells expressing aPKC D388A it was depolarized suggesting that aPKC D388A influences the localization or activity of endogenous aPKC (Figure 1C). We do not know the origin of the differential effects of aPKC K293W and aPKC D388A on Mira localization, but it may arise from differences in the amounts of the two proteins and how endogenous aPKC is affected.

We also examined the localization of the aPKC variants during interphase, when membrane-bound aPKC is normally not detectable, and found that the kinase inactivating mutations were predominantly found in the nucleus, presumably due to an embedded nuclear localization signal (Perander et al., 2001; Seidl et al., 2012). However, we also found that the aPKC kinase domains variants were enriched at the membrane relative to the cytoplasm (at sites away from progeny cell contacts) although at a level somewhat lower than apical WT protein at metaphase (Figure 1F,G).

Baz and Par-6 are normally found at the apical membrane with aPKC at metaphase. We examined whether the kinase inactive aPKC mutants influence Baz or Par-6 localization (Figure 2). We found that Baz remained apically polarized in cells expressing aPKC, as well as those expressing the aPKC D388A or aPKC K293W variants, although the intensity of this crescent was slightly reduced in aPKC D388A (Figure 2A, C). While Baz localization was unperturbed by the expression of the kinase inactive aPKC variants, Par-6 expanded into the basal domain like the localization of aPKC D388A and aPKC K293W (Figure 2A, B). We conclude that expression of kinase inactive aPKC leads to loss of Par-6 polarity but does not influence the localization of Baz. Depolarization of Par-6 but not Baz has also been observed when aPKC was chemically inhibited (Hannaford et al., 2019).

**Figure 2.**
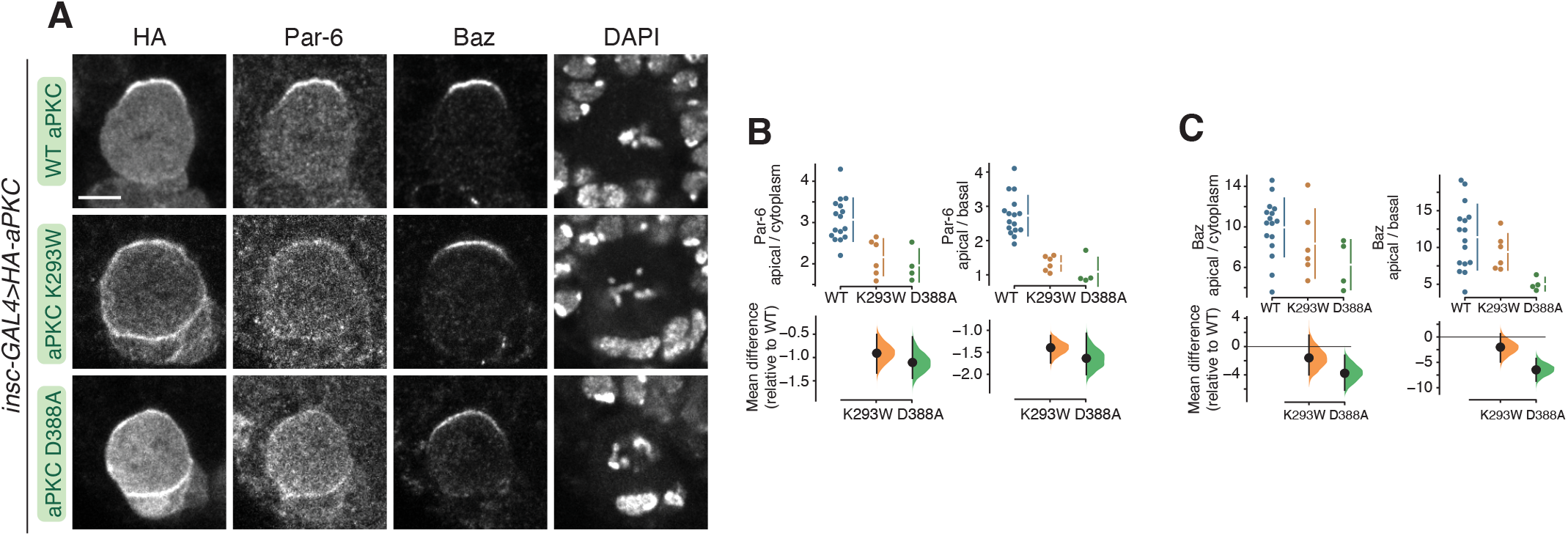
Localization of Bazooka and Par-6 in larval brain NSCs expressing kinase inactive aPKCs. **(A)** Localization of Par-6 and Bazooka (Baz) in metaphase larval brain NSCs expressing HA tagged aPKC D388A or aPKC K293W. Nucleic acids are shown with DAPI. Scale bar is 5 μm. **(B,C)** Gardner-Altman estimation plots of the effect of expressing aPKC D388A or aPKC K293W on Par-6 (B) or Baz (C) cortical localization and polarity. Apical cortical to cytoplasmic or basal cortical signal intensities of cortical and cytoplasmic signals are shown for individual metaphase NSCs expressing either HA-WT or HA-D388A or HA-K293W aPKC. Error bar in upper graphs represents one standard deviation (gap is mean); error bar in lower graphs represents bootstrap 95% confidence interval.

### Kinase inactive aPKC may bind the NSC membrane independently of Cdc42 and Bazooka

Membrane targeting of aPKC normally requires the activities of Baz and the small GTPase Cdc42 (Wodarz et al., 2000; Rolls et al., 2003; Atwood et al., 2007). We tested whether these upstream regulators are required for the uniformly cortical localization of kinase inactive aPKC by examining the localization of aPKC K293W in NSCs expressing Baz or Cdc42 RNAi. We found that wild-type aPKC failed to recruit to the membrane in these contexts, as previously reported (Figure 3A-D) (Atwood et al., 2007; Rolls et al., 2003). Consistent with this observation, we found that Mira was depolarized, localizing uniformly to the cortex in these cells. In contrast to wild-type aPKC, however, aPKC K293W was able to remain on the cortex even when Cdc42 was absent (Figure 3A-B). We also examined the localization of the aPKC K293W variant in NSCs expressing Baz RNAi. In this context, less aPKC is recruited to the apical membrane and Mira becomes depolarized as previously reported (Atwood et al., 2007). However, aPKC K293W’s remained highly enriched on the membrane (i.e., more than WT aPKC) when Baz was reduced (Figure 3C-D). While we cannot completely exclude a role for Cdc42 and Baz in recruiting aPKC K293W to the membrane, we conclude that aPKC K293W is targeted to the membrane significantly more than WT aPKC in metaphase NSCs with reduced Cdc42 or Baz function.

**Figure 3.**
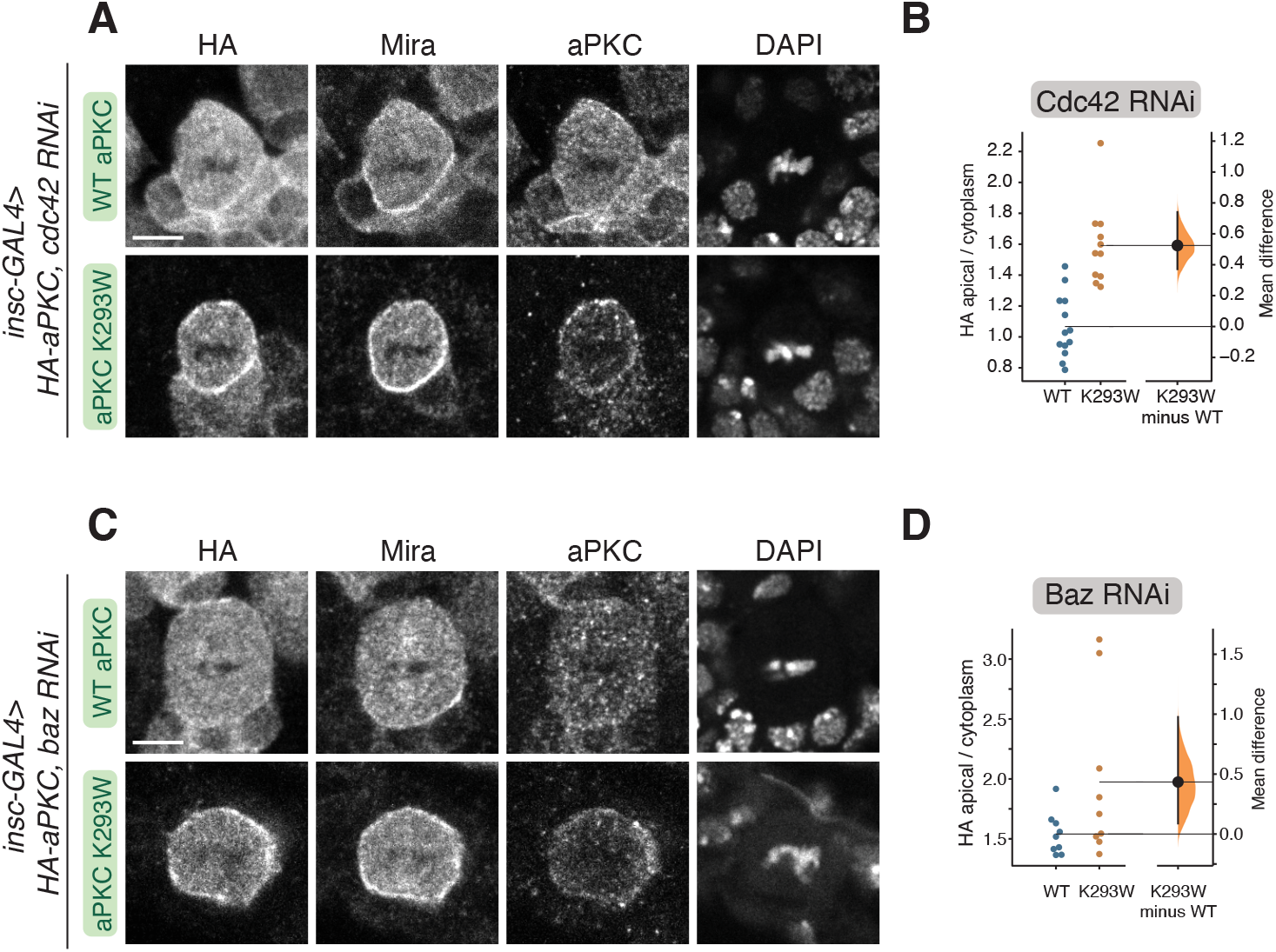
Cortical localization of kinase inactive aPKC in NSCs lacking Bazooka or Cdc42. **(A)** Localization of HA tagged aPKC K293W in metaphase larval brain NSCs expressing an RNAi directed against Cdc42. Scale bar is 5 μm in all panels. **(B)** Gardner-Altman estimation plots of the effect of expressing Cdc42 RNAi on WT and K293W aPKC cortical localization. Apical cortical to cytoplasmic signal intensities of anti-HA signals are shown for individual metaphase NSCs expressing either HA-WT or HA-K293W aPKC. Error bar represents bootstrap 95% confidence interval. **(C)** Localization of HA tagged aPKC K293W in metaphase larval brain NSCs expressing an RNAi directed against Bazooka. **(D)** Gardner-Altman estimation plots of the effect of expressing Baz RNAi on WT and K293W aPKC cortical localization. Apical cortical to cytoplasmic signal intensities of anti-HA signals are shown for individual metaphase NSCs expressing either HA-WT or HA-K293W aPKC. Error bar represents bootstrap 95% confidence interval.

### The aPKC C1 domain is a constitutive membrane targeting module

Our results indicate that the uniform membrane localization of aPKC with inactive kinase domains (e.g., K293W) is independent of both Baz and Cdc42. One model consistent with these results is that aPKC contains a membrane targeting module that is regulated by its kinase domain, along with Baz and Cdc42 binding. To identify the putative module, we first examined whether removing the kinase domain leads to the same uniform membrane localization phenotype. We found that aPKC PB1-C1 (aka ΔKD; Figure 4A) was enriched at the membrane and not polarized (Figure 4B-D). A similar localization pattern has been reported for aPKC PB1-C1 expressed in cultured cells (Dong et al., 2020). Thus, aPKC’s NH2-terminal regulatory region, consisting of PB1, PS, and C1 domains, is responsible for membrane binding. The PB1 could mediate interaction with the membrane via protein-protein interactions with Par-6 (Atwood et al., 2007; Petronczki and Knoblich, 2001), the PS domain through direct interactions with lipids (Dong et al., 2020), or the C1 domain which serves as a membrane targeting module in other PKCs (Colón-González and Kazanietz, 2006) but has not been reported to do so in aPKCs. Removal of the C1 domain from the regulatory domain (aPKC PB1-PS) leaving the PB1 and PS domains (Figure 4A), resulted in a protein that was not enriched at the membrane (Figure 4B-D). Thus, the PB1 and PS domains are not sufficient for membrane targeting in NSCs. We next tested the C1 domain alone and found that it was enriched uniformly to the membrane relative to the cytoplasm (Figure 4B-D). We also observed C1 enrichment at the membrane during interphase (Figure 4E), although the domain was predominantly found in the nucleus, presumably due to an embedded nuclear localization signal (Perander et al., 2001; Seidl et al., 2012). We conclude that the aPKC C1 domain is a membrane targeting module.

**Figure 4.**
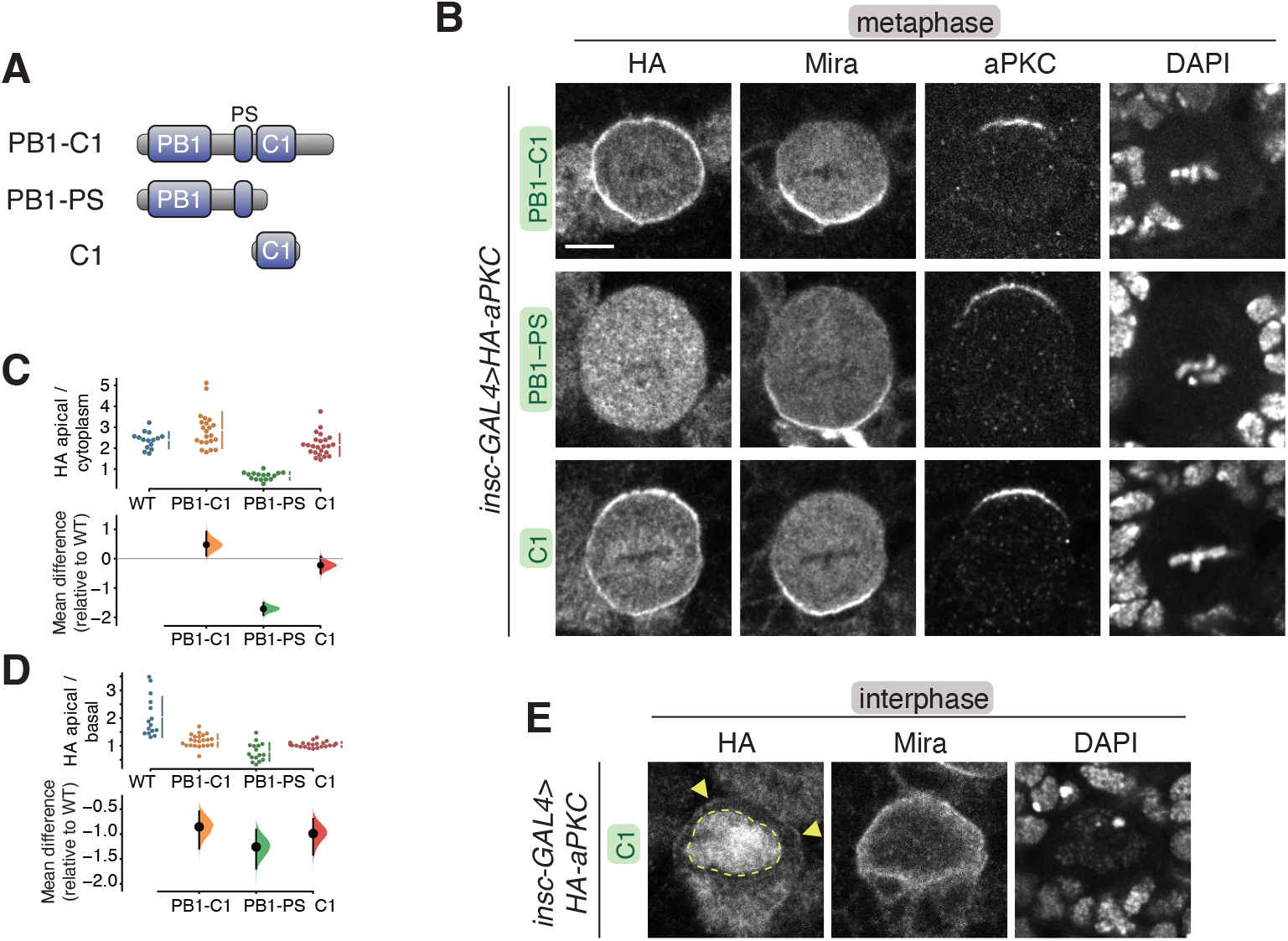
Localization of the aPKC regulatory domain in larval brain NSCs. **(A)** aPKC regulatory domain fragments. **(B)** Localization of HA tagged aPKC regulatory domain fragments in metaphase larval brain NSCs. The basal marker Miranda, endogenous aPKC (using an antibody that does not react with the regulatory domain), nucleic acids (DAPI) are shown for comparison. Scale bar is 5 μm. **(C,D)** Gardner-Altman estimation plot of aPKC regulatory domain cortical localization (C) and polarity (D). Apical cortical to cytoplasmic (C) and apical cortical to basal cortical signal intensity ratios (D) of anti-HA signals are shown for individual metaphase NSCs expressing either aPKC PB1-C1, PB1-PS, or C1 regulatory domain fragments. Error bar in upper graphs represents one standard deviation (gap is mean); error bar in lower graphs represents bootstrap 95% confidence interval. **(E)** Localization of the HA tagged aPKC C1 domain in interphase larval brain NSCs. Arrowheads highlight the membrane signal and the nuclear signal is outlined by a dashed line.

### C1 is a lipid binding module that is required for aPKC membrane recruitment

How might the C1 domain mediate interaction with the NSC membrane? C1 domains from canonical PKCs bind diacylglycerol (DAG), although the aPKC C1 domain does not bind DAG (Colón-González and Kazanietz, 2006). However, we sought to determine if aPKC’s C1 domain binds other phospholipids. We used a vesicle pelleting assay in which Giant Unilamellar Vesicles (GUVs) with varying phospholipid compositions were mixed with purified aPKC C1 domain. The vesicles were separated from the soluble phase by ultracentrifugation and any associated C1 was identified by protein gel electrophoresis. We observed varying degrees of C1 binding to a broad array of phospholipids (Figure 5A), suggesting that the C1 is a nonspecific phospholipid binding module.

**Figure 5.**
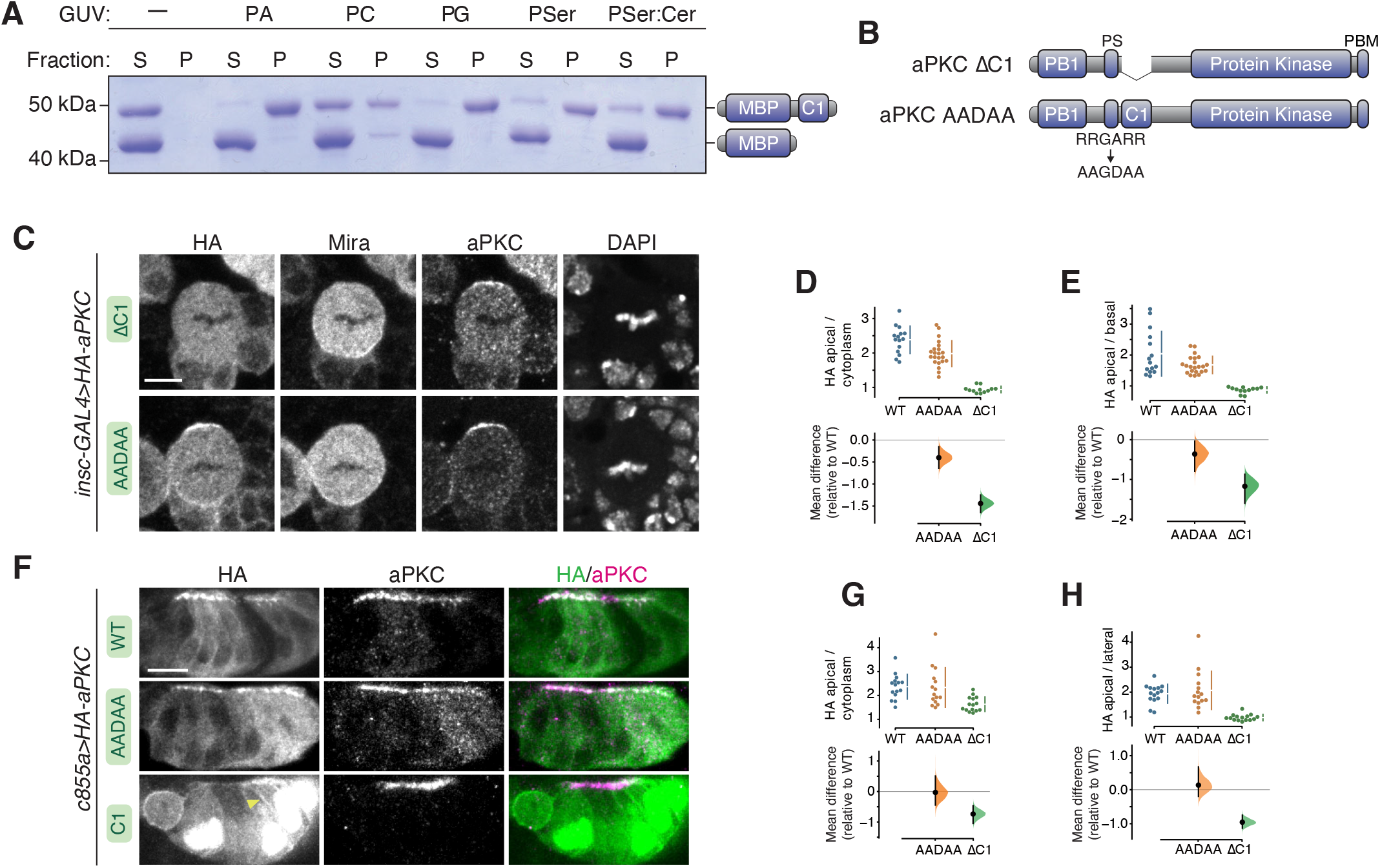
Phospholipid binding of aPKC C1 domain and role of C1 and PS domains in aPKC localization in larval brain NSCs and epithelia. **(A)** Binding of a maltose binding protein (MBP) fusion of the aPKC C1 domain to phospholipids. Supernatant (S) and pellet (P) fractions from cosedimentation with Giant Unilamellar Vesicles (GUVs) of the indicated phospholipid composition are shown (PA, phosphatidic acid; PC, phosphatidyl choline; PG, phosphatidyl glycerol; PSer, phosphatidyl serine; PSer:Cer, phosphatidyl serine mixture with ceramide). MBP alone is included as an internal negative control. **(B)** Schematics of ΔC1 and AADAA aPKC variants. **(C)** Localization of HA tagged aPKC ΔC1 and AADAA variants in metaphase larval brain NSCs. The basal marker Miranda, total aPKC (expressed variant and endogenous), nucleic acids (DAPI) are shown for comparison. Scale bar is 5 μm. **(D,E)** Gardner-Altman estimation plots of aPKC AADAA and ΔC1 cortical localization in NSCs. Apical cortical to cytoplasmic (D) or apical to basal (E) signal intensity ratios of anti-HA signals are shown for individual metaphase NSCs expressing either aPKC AADAA or ΔC1. Error bar in upper graphs represents one standard deviation (gap is mean); error bar in lower graphs represents bootstrap 95% confidence interval. **(F)** Localization of HA tagged aPKC ΔC1 and AADAA variants in larval brain inner proliferation center (IPC) epithelium. Arrowhead highlights aPKC C1 localization at the lateral membrane. As in interphase NSC cells, the C1 is highly enriched in the epithelial nuclei. Scale bar is 5 μm. **(G,H)** Gardner-Altman estimation plots of aPKC AADAA and C1 cortical localization in IPC epithelial cells. Apical cortical to cytoplasmic (D) or apical to lateral (E) signal intensity ratios of anti-HA signals are shown for individual epithelial cells from the IPC expressing either aPKC AADAA or C1. Error bar in upper graphs represents one standard deviation (gap is mean); error bar in lower graphs represents bootstrap 95% confidence interval.

To better understand the role of the C1 in aPKC polarity, we examined the effect of removing it (aPKC ΔC1; Figure 5B) on aPKC’s localization. We found that aPKC ΔC1 remained in the cytoplasm and was not enriched at the membrane (Figure 5C-E), leading us to conclude that the C1 is required for aPKC membrane targeting in NSCs. Interestingly, Mira localization was also disrupted in NSCs expressing aPKC ΔC1 suggesting that the C1 also plays a role in regulating aPKC’s protein kinase activity in NSCs. The displacement of cortical Mira in NSCs expressing aPKC ΔC1 is consistent with the increase in catalytic activity observed in *in vitro* measurements of this protein (Graybill et al., 2012; Zhang et al., 2014).

The PS domain has been reported to be a membrane binding module required for membrane recruitment of aPKC in cultured cells and epithelia (Dong et al., 2020). Our results suggest that the PS is not sufficient for localization to the NSC membrane as aPKC PB1-PS remains in the cytoplasm. We tested whether the PS is required for NSC aPKC membrane recruitment by examining the localization of aPKC in which the positively charged, basic residues were removed and a negatively charged side chain added (aPKC AADAA; Figure 5B). This mutation has been reported to abrogate membrane binding in contexts where the PS is required (Dong et al., 2020). We found that the apical localization of aPKC AADAA in NSCs was slightly reduced compared to WT aPKC, suggesting that the PS contributes to the recruitment process but is not absolutely required for recruitment or polarization in this cellular context (Figure 5C-E). Mira localization was cytoplasmic in aPKC AADAA-expressing NSCs, however, consistent with the increase in kinase activity that is expected when the autoinhibitory PS is inactivated (Graybill et al., 2012). We also assessed the localization of aPKC AADAA in the epithelium of the larval brain inner proliferation center by expressing it with the c855a-GAL4 driver. We found this protein to be enriched at the apical membrane of interphase epithelial cells in a similar pattern to wild-type aPKC (Figure 5F-H). We also examined the localization of the isolated C1 in interphase epithelial cells and observed a similar pattern to NSCs – predominantly nuclear but also enriched at the membrane relative to the cytoplasm (Figure 5F-H).

### Regulation of membrane recruitment by the PB1 domain and its interaction with Par-6

Our results indicate that recruitment of aPKC to the membrane is mediated primarily by the C1 domain. The constitutive membrane binding of the isolated C1 suggests that other aPKC domains regulate the C1 through autoinhibition to yield the dynamic membrane localization of full-length aPKC. In this model, perturbation of a regulatory domain could lead to constitutively cytoplasmic or membrane-bound aPKC if the perturbation disrupted C1 activation or repression, respectively. Cytoplasmic aPKC has been observed upon inactivation of aPKC’s PDZ Binding Motif (PBM; Holly et al., 2020), and kinase domain active site perturbations lead to uniform membrane binding (Figure 1) (Hannaford et al., 2019; Rodriguez et al., 2017). We determined if aPKC’s PB1 domain (Figure 1A) participates in C1 regulation by examining the localization of aPKC D77A, a PB1 point mutation that disrupts interaction with Par-6’s PB1 domain (Hirano et al., 2004), and aPKC ΔPB1, which lacks the PB1 entirely (Figure 6A). Intriguingly, each perturbation leads to a distinct effect – disrupting the interaction with Par-6 leads to cytoplasmic aPKC but complete deletion of the PB1 leads to constitutive membrane binding (Figure 6B-D). The distinct localization resulting from these two perturbations indicate that the PB1 is required to repress aPKC membrane recruitment and PB1’s interaction with Par-6 is required to overcome this regulation.

**Figure 6.**
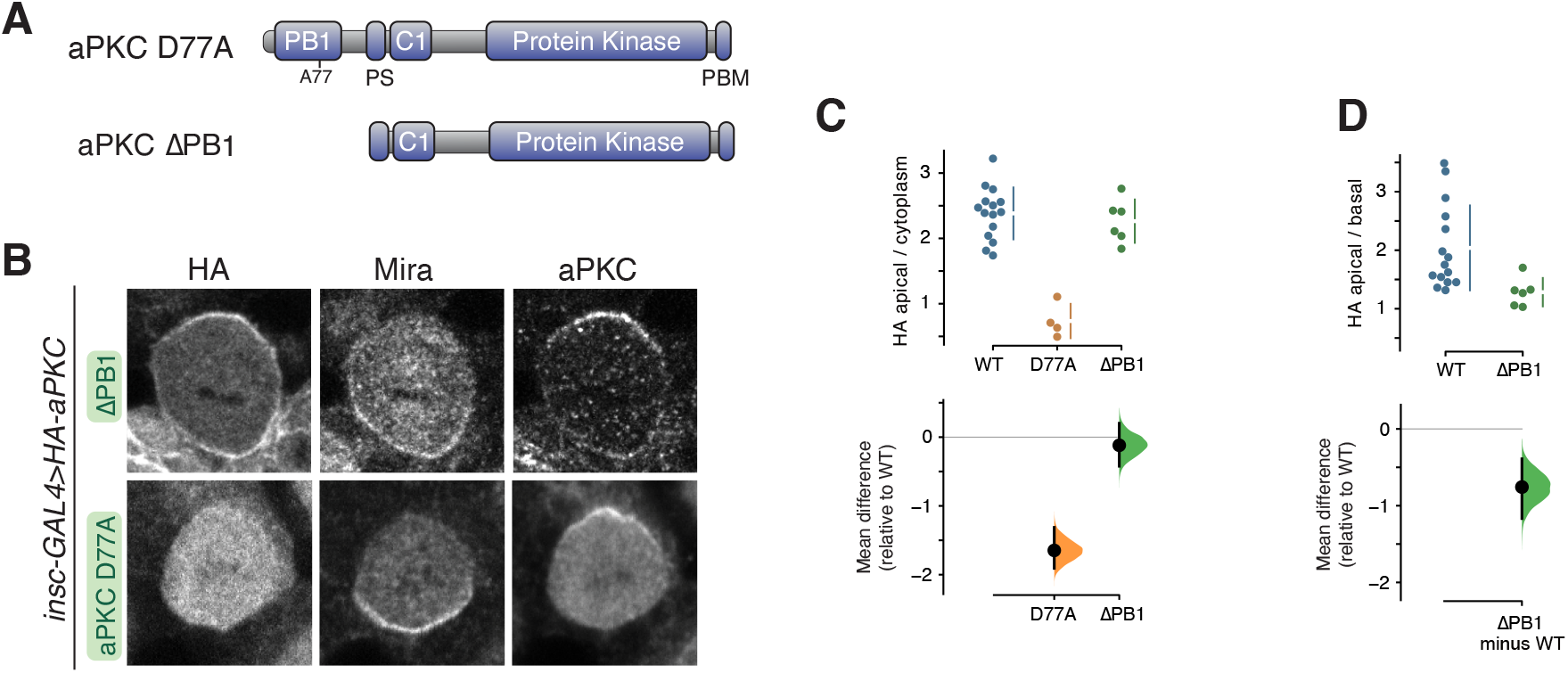
Localization of aPKC with PB1 domain perturbations in larval brain NSCs. **(A)** Schematics of D77A and ΔPB1 aPKC variants. **(B)** Localization of HA tagged aPKC D77A and ΔPB1 variants in metaphase larval brain NSCs. The basal marker Miranda, and total aPKC (expressed variant and endogenous), are shown for comparison. Scale bar is 5 μm. **(C,D)** Gardner-Altman estimation plots of aPKC D77A and ΔPB1 cortical localization. Apical cortical to cytoplasmic (C) or apical to basal (D) signal intensity ratios of anti-HA signals are shown for individual metaphase NSCs expressing either aPKC D77A or ΔPB1. Error bar in upper graphs represents one standard deviation (gap is mean); error bar in lower graphs represents bootstrap 95% confidence interval.

## Discussion

The Par complex component aPKC undergoes a dynamic localization cycle in NSCs, targeting to the apical membrane briefly in mitosis and returning to the cytoplasm as division completes (Oon and Prehoda, 2019). The function of interphase, cytoplasmic localization is unknown, but the apical localization of aPKC during mitosis is necessary for the polarization of fate determinants (Atwood et al., 2007; Prehoda, 2009; Rolls et al., 2003), a prerequisite for asymmetric cell division. Regulated membrane recruitment of aPKC is a central aspect of Par-mediated polarity and many physical interactions between aPKC and proteins and phospholipids have been identified. Conceptually, targeting could occur through the concerted action of multiple weak interactions (i.e., avidity). However, we discovered that the aPKC C1 domain is a phospholipid binding module sufficient for membrane recruitment, whereas domains that mediate protein-protein interactions or other interactions with phospholipids are not sufficient for aPKC targeting. As the C1 is the only domain within aPKC with this capability in NSCs, our results suggest that other aPKC domains, including the catalytic domain, function, at least in part, to regulate the membrane recruitment activity of the C1. While the PS and C1 domains are both known to form intramolecular interactions with the catalytic domain (Graybill et al., 2012; Zhang et al., 2014), a predicted structure of aPKC suggests that there are also significant intramolecular interactions within the regulatory PB1-PS-C1 module (Figure 7A) (Jumper et al., 2021; Varadi et al., 2022). Taken collectively, we propose that cooperative activation of the C1 leads to the spatially and temporally controlled localization of aPKC observed in many animal cells.

**Figure 7.**
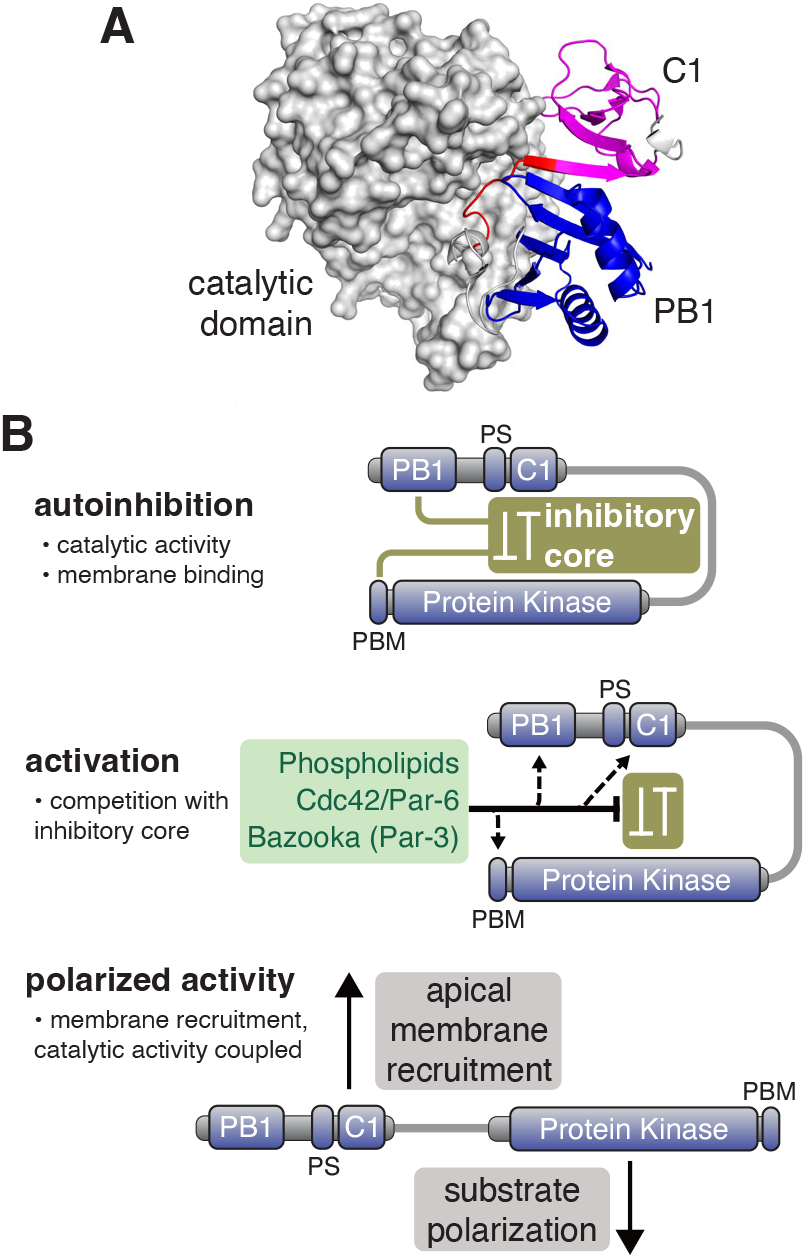
Model for regulation of aPKC activity and membrane recruitment. **(A)** Alphafold database structure (UNIPROT: A1Z9X0) of aPKC showing putative supramolecular architecture of PB1 (blue), PS (red) and C1 (magenta) domains and interaction of both domains with the catalytic domain. **(B)** Model for cooperative polarization and activation of aPKC. An inhibitory core couples repression of catalytic activity (protein kinase domain) to membrane localization (C1 with some contribtion from PS). The PB1 and PBM are also coupled to the inhibitory core to allow for cooperative activation by Cdc42/Par-6 binding to the PB1 and Baz binding to the PBM. Disruption of the inhibitory core leads to spatially (apical) and temporally (mitotic) regulated localization and activation of catalytic activity.

The C1 is unique in its ability to promote aPKC membrane recruitment in NSCs. While numerous interactions between phospholipids and domains outside the C1 have been reported (Wang et al., 2005; Ivey et al., 2014; Kajimoto et al., 2019; Dong et al., 2020), our results indicate that they are not sufficient for membrane recruitment in NSCs. Similarly, the domains that mediate protein-protein interactions, such as the PB1 that binds Par-6 and the PBM that binds Baz, are also not sufficient for aPKC recruitment. While the isolated Par-6 and aPKC PB1 domains bind one another *in vitro*, our data indicate that the aPKC PB1 domain is not sufficient for membrane targeting, perhaps because Par-6 itself requires aPKC for membrane targeting (Rolls et al., 2003).

The constitutive nature of C1 membrane binding raises the question of how its activity is regulated to yield aPKC’s precise spatially and temporally controlled localization. The aPKC catalytic domain forms intramolecular interactions with the PS and C1 that repress kinase activity (Graybill et al., 2012; Zhang et al., 2014), forming an “inhibitory core” (Figure 7B). Our results suggest the inhibitory core is coupled to C1 membrane binding. We observed C1 localization at the membrane throughout the cell cycle and uniform membrane binding in mitosis when aPKC is normally restricted to the apical hemisphere. We also observed constitutive membrane recruitment in aPKC variants where the catalytic domain was perturbed, suggesting that it is required for repression of C1 activity. We suggest that perturbations to the catalytic domain that influence protein kinase activity can also disrupt the inhibitory core, consistent with the complex allosteric pathways in eukaryotic protein kinase domains (Ahuja et al., 2019). It has been proposed that the PS plays a central role in membrane recruitment and coupling localization to the inhibitory core through its interactions with the catalytic domain (Dong et al., 2020). Our results indicate that the PS is not sufficient for membrane recruitment, nor is its interaction with the aPKC catalytic domain required for polarization in NSCs or in a larval epithelium – aPKC AADAA is polarized but constitutively active. We suggest that the PS plays a more significant role in regulating catalytic activity than localization in these tissues.

Given that the C1 appears to be autoinhibited by the catalytic domain, how might it become activated? We previously found that inactivation of the aPKC PBM (aPKC V606A), which binds Baz, leads to cytoplasmic aPKC localization (Holly et al., 2020). Taken with our current results, we suggest that the interaction with Baz is required for membrane recruitment not because Baz directly recruits aPKC (e.g., aPKC ΔC1 is cytoplasmic), but because the Baz interaction is required for disruption of the inhibitory core and activation of the C1. Similarly, in NSCs lacking the PB1-binding Par-6 protein, aPKC also remains in the cytoplasm, even though Baz remains properly polarized and could potentially bind aPKC’s PBM (Rolls et al., 2003). These observations underlie our emphasis on the cooperative nature of aPKC membrane recruitment – activation of the C1 by at least Par-6 and Baz leads to the complex localization dynamics of aPKC observed in NSCs. Future work will be directed at understanding how the aPKC PB1 and PBM might be coupled to the inhibitory core and activation of aPKC membrane binding.

## Materials and Methods

### Drosophila

Flies were grown at indicated temperatures on standard cornmeal/yeast media. Both male and female larvae were used in this study. Transgenic constructs were cloned into the pUAST attB vector (GenBank: EF362409.1) modified to include an N-terminal 3xHA or 1xHA tag. Integration of the vectors was done using standard Phi-C31 integration into an attP landing site on the third chromosome (attP2) by Rainbow Genetics or BestGene Inc. Positive insertion was determined by the presence of colored eyes after backcrossing to y,w stock.

### Immunofluorescence

For analysis of neural stem cells, *insc*-Gal4 virgins were crossed to males containing an aPKC transgene on the third chromosome or an RNAi on the second chromosome and an aPKC transgene on the third chromosome. Crosses laid in vials for 24 hours at ~20 °C. The resulting embryos were incubated at 30 °C until larvae reached third instar wandering larva stage. For optic lobe experiments, *c855a*-Gal4 virgins were crossed to males containing an aPKC transgene on the third chromosome. Crosses laid in vials for 7 days (~20 °C) and the resulting progeny were incubated at 30 °C for 24 hours. Wandering third instar larva were selected for dissection. Following dissection, the tissue was incubated in 4% PFA fixative for 20’ within 20’ of dissection. This and all subsequent wash steps involved agitation by placing on a nutator. After fixation, brains were rinsed and washed three times for 15’ each in PBST (1xPBS with 0.3% Triton-X). Brains were stored for up to three days at 4 °C before staining. Before staining, brains were blocked for 30’ in PBSBT (PBST with 1% BSA) and then incubated with primary antibody solution overnight at 4 °C. Brains were subsequently rinsed and washed three times for 15’ each in PBSBT and incubated with secondary antibody for 2 hours in a vessel that protected the sample from light. Brains were subsequently rinsed and washed three times for 15’ each in PBST followed by storage in SlowFade w/DAPI at least overnight before imaging. Brains were imaged on an upright Leica TCS SPE confocal using an ACS APO 40x 1.15 NA Oil CS objective.

### MARCM Immunofluorescence

To create MARCM larval NSC clones, FRT-G13, aPKC^K06403^/CyO virgins were crossed with 3xHA aPKC D388A males. The subsequent progeny were allowed to grow to adulthood and were screened for the absence of the CyO marker. Males with no CyO were then crossed to elav-Gal4, UAS-mCD8:GFP, hs:flp; FRT-G13, tubPGal80 virgins.

Crosses laid in vials for 24 hours and the resulting embryos were incubated at room temperature (~20 °C) for 24 hours. These vials were then heat shocked at 37 °C for 90 min. Another heat shock was possible within 18 hours. Larvae were allowed to grow at room temperature or 18 °C until third instar wandering larva stage, when they were dissected and fixed as above.

### Membrane enrichment and polarization quantification

A 10 pixel-wide line from apical to basal membrane in the medial optical section was used to measure membrane signals in metaphase NSCs. The signal at the edge of the cell was used as the membrane signal and the cytoplasmic signal was taken as the average of 20 data points located 10 points from the apical peak.

For interphase NSCs and epithelial cells, cortical intensity was measured by tracing the cortex with a 3px line, while cytoplasmic signal was taken as an average of the entire cytoplasmic signal.

All images were analyzed using Fiji and statistical analysis was done using DABEST python package (https://github.com/ACCLAB/DABEST-python).

Figures were assembled using Adobe Illustrator.

### Vesicle cosedimentation assay

MBP-C1 was purified use amylose agarose affinity purification as previously described (Graybill et al., 2012). For Giant Unilamellar Vesical (GUV) production, 50 μl of the specified lipids at 10mg/mL in chloroform was dried in a test tube under an N_2_ stream and then in a vacuum chamber to ensure all chloroform was removed. Lipids were resuspended in a 0.2M sucrose solution to a final concentration of 0.5mg/mL and heated in a water bath at 50 °C for 5 hours with occasional agitation. All lipids were stored at 4 °C and used within 3 days. All spins were carried out using an Optima MAX-TL Ultracentrifuge with a TLA-100 rotor at 65,000 at 4 °C.

MBP-C1 protein was diluted to 50μM in 20 mM HEPES, pH 7.5; 50 mM NaCl, 1 mM DTT and pre-cleared for 30’. Reaction conditions: 20mM HEPES, pH 7.5, 50mM NaCl, 1mM DTT, 0.25mg/mL GUVs, and 5μM MBP-C1. Reaction was carried out at room temperature for 15’ and then spun down for 30’. The supernatant fraction was removed, and the pellet was resuspended in an equivalent volume of 1X Dilution buffer. Both the supernatant and pellet samples were mixed with 6x loading dye and separated using 12.5% SDS-PAGE. Gels were stained with Coomassie and imaged using a scanner.

## Key Resources Table

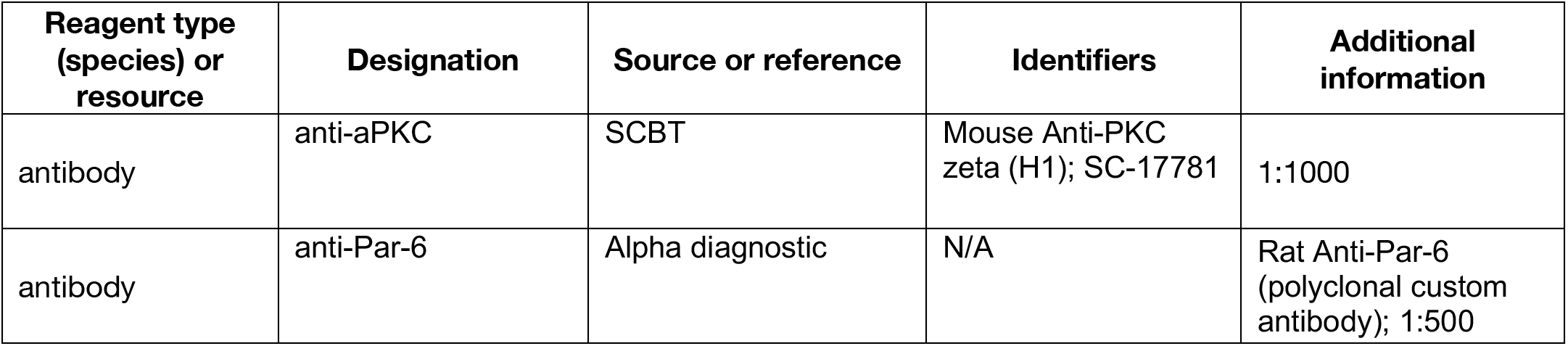
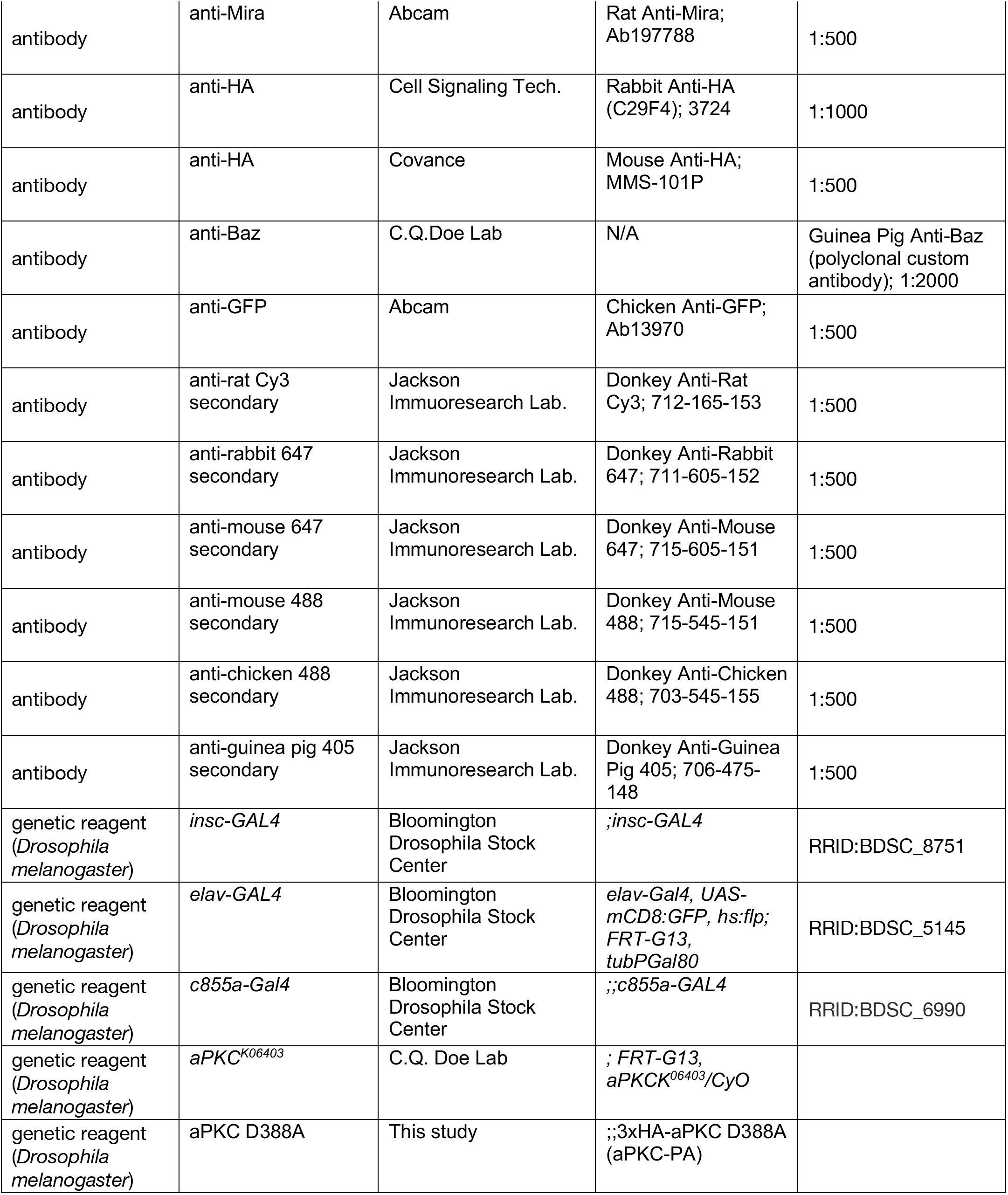
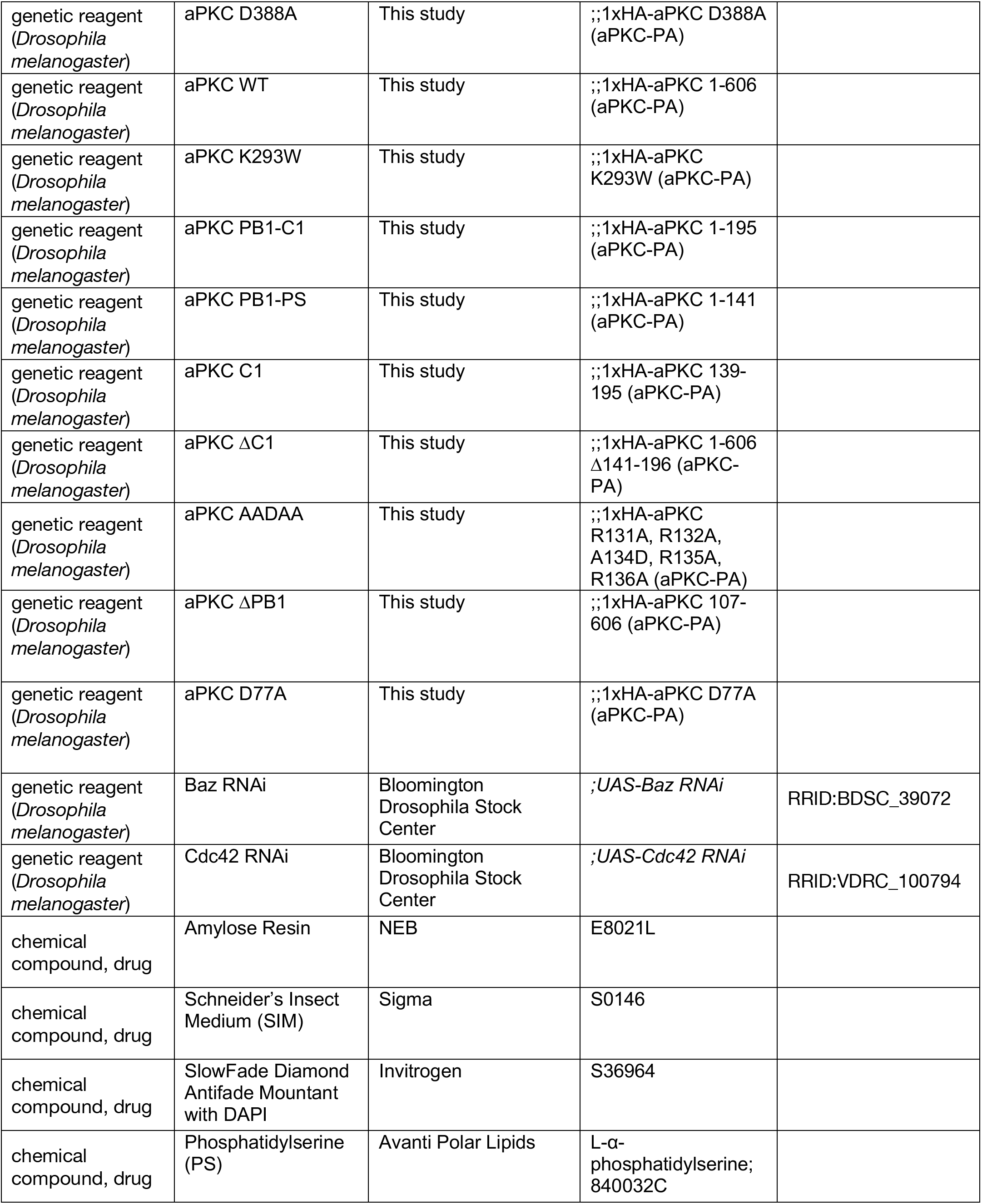
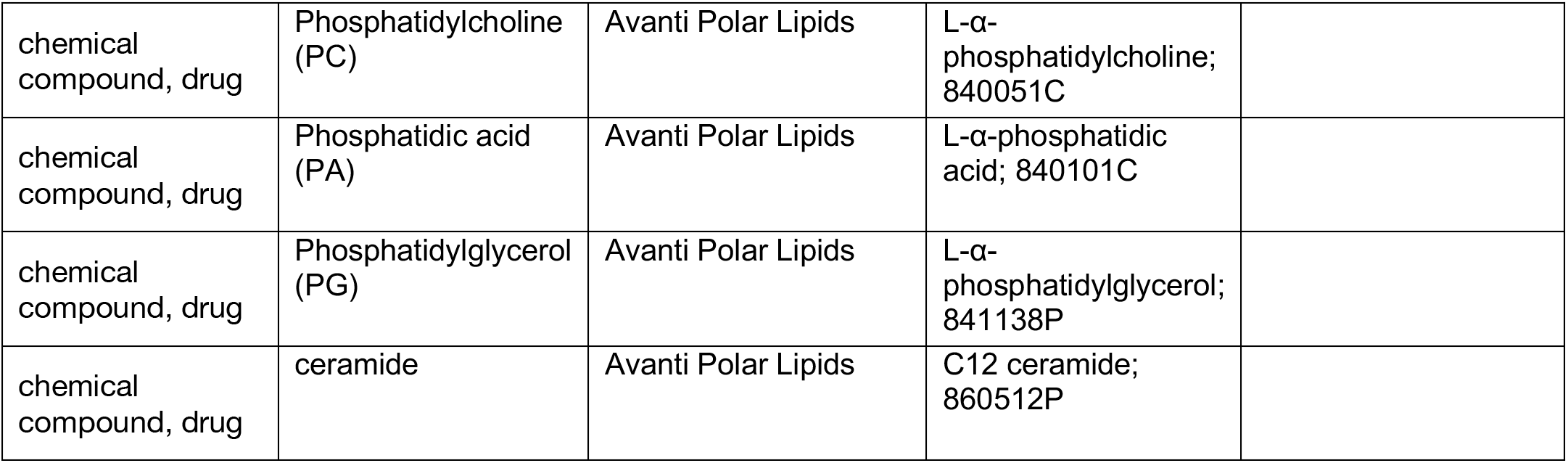

